# CaSpER: Identification, visualization and integrative analysis of CNV events in multiscale resolution using single-cell or bulk RNA sequencing data

**DOI:** 10.1101/426122

**Authors:** Akdes Serin Harmancı, Arif O. Harmanci, Xiaobo Zhou

## Abstract

RNA sequencing experiments generate large amounts of information about expression levels of genes. Although they are mainly used for quantifying expression levels, they contain much more biologically important information such as copy number variants (CNV). Here, we propose CaSpER, a signal processing approach for identification, visualization, and integrative analysis of focal and large-scale CNV events in multiscale resolution using either bulk or single-cell RNA sequencing data. CaSpER performs smoothing of the genome-wide RNA sequencing signal profiles in different multiscale resolutions, identifying CNV events at different length scales. CaSpER also employs a novel methodology for generation of genome-wide B-allele frequency (BAF) signal profile from the reads and utilizes it in multiscale fashion for correction of CNV calls. The shift in allelic signal is used to quantify the loss-of-heterozygosity (LOH) which is valuable for CNV identification. CaSpER uses Hidden Markov Models (HMM) to assign copy number states to regions. The multiscale nature of CaSpER enables comprehensive analysis of focal and large-scale CNVs and LOH segments. CaSpER performs well in accuracy compared to gold standard SNP genotyping arrays. In particular, analysis of single cell Glioblastoma (GBM) RNA sequencing data with CaSpER reveals novel mutually exclusive and co-occurring CNV sub-clones at different length scales. Moreover, CaSpER discovers gene expression signatures of CNV sub-clones, performs gene ontology (GO) enrichment analysis and identifies potential therapeutic targets for the sub-clones. CaSpER increases the utility of RNA-sequencing datasets and complements other tools for complete characterization and visualization of the genomic and transcriptomic landscape of single cell and bulk RNA sequencing data, especially in cancer research.

## Introduction

Tumors are complex ecosystems composed of heterogeneous cell populations^1^. Understanding the clonal cellular composition of the tumor and the interplay between the cells within the tumor ecosystem provides significant insights in the tumor recurrence, treatment, initiation, progression, and metastasis^2^. With the transformative advances in experimental methods, such as next-generation sequencing methods, we are now able to build the complex map of the tumor ecosystem utilizing the deeply sequenced transcriptome and the genome of the cells within the tissue.

Over the past few years, the development and application of single-cell DNA and RNA sequencing methods have revolutionized cancer research^3^. Single cell RNA-sequencing (scRNA-Seq) is a powerful new deep molecular profiling method for detecting different cell types, states and functions in cancer^3–10^. Several previous studies characterized the heterogeneity and crosstalk within tumor microenvironment in various cancer types using scRNA-Seq data^4,6,8^. Single-cell DNA sequencing is another new powerful approach for understanding the genomic diversity of tumor clonal architecture^3,11^. However, it remains technically challenging to assay both the genome and transcriptome from the same cell.

RNA-sequencing experiments are predominantly performed for the purpose of estimating gene activity through quantification of gene and transcript. These datasets, however, contain a substantial amount of information about the genomic variants in the samples and are severely underutilized. For example, RNA-Seq data has been used to identify single nucleotide polymorphisms (SNP) and short indels^12–14^. Identification of these variants from RNA-Seq data increases the utility of RNA-Seq experiments significantly compared to using RNA-Seq only for gene expression quantification because researchers can integrate a portion of the genomic landscape of the tumor cells (as much as it is revealed by RNA-Seq) with the transcriptomic landscape rather than studying the transcriptomic landscape of the cells alone. Identification of other variants can enable an even more complete characterization and present higher utility for RNA-Seq data. Among these variants, copy number variants (CNVs) are very important for cancer research because they are a major class of genetic drivers of cancer. Complete losses and gains of genomic material can cause loss and gain of tumor-suppressor and oncogenes, respectively and cause transformation of healthy cells into tumor cells. Inference of copy number variation (CNV) in the tumor samples is essential for understanding the correlation between the genomic and the transcriptomic properties of different cell types and clones within the tumor ecosystem^15^. These correlations will provide significant insight into the tumor initiation, progression, and metastasis. Identification of CNVs from RNA-Seq data, however, is very challenging because the dynamic and highly non-uniform coverage of the genome by RNA-Seq signal makes it very hard to distinguish between deletions/amplifications and dynamic variation of gene expression levels. Considering the growing number of RNA-Sequencing studies, especially with the release of TCGA^16^, ENCODE^17^, GTEx^18^, Human Cell Atlas (HCA)^19^, Human Tumor Atlas Network (HTAN), and Human Biomolecular Atlas Program (HuBMAP) consortium datasets, there is an increasing need for developing CNV inference algorithm from RNA-Seq data. New algorithms can substantially increase the utility of these existing datasets.

In this paper, we focus on systematic identification of CNVs using bulk and single cell RNA-sequencing datasets. We present CaSpER, a statistical framework for analysis and visualization of the genomewide RNA-Seq signal profiles. Many algorithms have been developed for detecting CNV events from DNA sequencing data using depth of coverage analysis^20,21^. These tools rely on uniform coverage of genome by DNA-sequencing reads. However, statistical approaches for CNV detection using RNA-sequencing data is very limited since it is very hard to discriminate between differential expression and an underlying copy number variation using only RNA-Seq data. Another challenge is that RNA-Seq signal is generally concentrated on the exonic regions and most of the genome is not covered. Thus, the identified CNVs will reflect the copy number states of the genes and the copy number of intergenic regions may not be represented well. Regardless, the copy number of genes is extremely useful information for characterizing CNV architecture of, for example, the copy number of oncogenes and tumor suppressor genes. It is worth noting that this issue is similar to the whole exome-sequencing based CNV detection because the whole exome-sequencing covers only the targeted exonic regions in the genome.

Although there are many tools that identify CNVs from exome sequencing data, there is much scarcity of methods for detecting CNVs solely from RNA sequencing data^4,22^. One relevant method is inferCNV^4^ which enables only visual inspection of expression profiles from scRNA-Seq, another method is HoneyBADGER^22^ which enables calling CNV from scRNA-Seq data.

One aspect that is very important and often overlooked is coherent identification, visualization, and integrative analysis of focal and large-scale CNV changes. To study CNV events at multiple scales, CaSpER utilizes a computational approach for identifying CNVs using a multiscale signal-processing framework. To increase the specificity of the identified CNVs, CaSpER utilizes a novel and efficient method to generate allelic shift signal profile. This profile quantifies the genome-wide loss-of-heterozygosity which has been previously shown to be extremely useful for identifying CNVs. Unlike most other tools, CaSpER does not require heterozygous variant calls to generate the allelic shift profile. CaSpER integrates the allelic shift profile in the multiscale analysis and assigns CNVs using Hidden Markov Models (HMM). CaSpER identifies and visualizes mutually exclusive and co-occurring CNV alterations and infers CNV based clonal evolution of tumor using single-cell RNA-Seq data. CaSpER also identifies gene expression signatures of mutually exclusive CNV sub-clones and performs gene ontology (GO) enrichment analysis. CaSpER enables visualization and integrative analysis of a large number of single-cell RNA-Seq datasets for a complete characterization of RNA-Seq data in large studies. We evaluate the accuracy of CaSpER and present use cases of CaSpER for both single-cell and bulk RNA-Sequencing data. Overall CaSpER complements the existing arsenal of cancer genome sequencing analysis tools for a complete understanding of the tumor architecture.

## Results

### CaSpER identifies CNV events from bulk or single-cell RNA-Sequencing data

The overview of CaSpER algorithm is shown in Figure 1. CaSpER uses expression values and B-allele frequencies (BAF) from RNA-Seq reads to estimate CNV events. The BAF is a relative normalized measure of the allelic intensity ratio of two alleles (A and B). The allele A is the reference allele whereas the allele B is the non-reference allele. The BAF value of 1 and 0 corresponds to absence of one allele, BB and AA consecutively, and the BAF value of 0.5 corresponds to presence of both alleles, AB. The input to CaSpER consists of aligned RNA-Seq reads and the window lengths to be used in multiscale analysis^23^. CaSpER first generates an expression signal by quantifying expression values of all the genes from aligned RNA-Seq reads. The expression values for the genes are treated as a genome-wide signal profile. In order to eliminate the noise in the initial expression signal profile, CaSpER performs sliding window based median filtering and computes the multiscale decomposition of the expression signal in multiple window length scales. The window length is increased between consecutive scales. Next, for the smoothed signal at each scale, a 5-CNV-state Hidden Markov Models (HMM) is used to assign copy number states to regions and segment the signal into regions of similar copy number states. We observed that pooling the samples in our bulk or single-cell RNA-Seq data gives better initial HMM parameter estimates.

**Figure 1.**
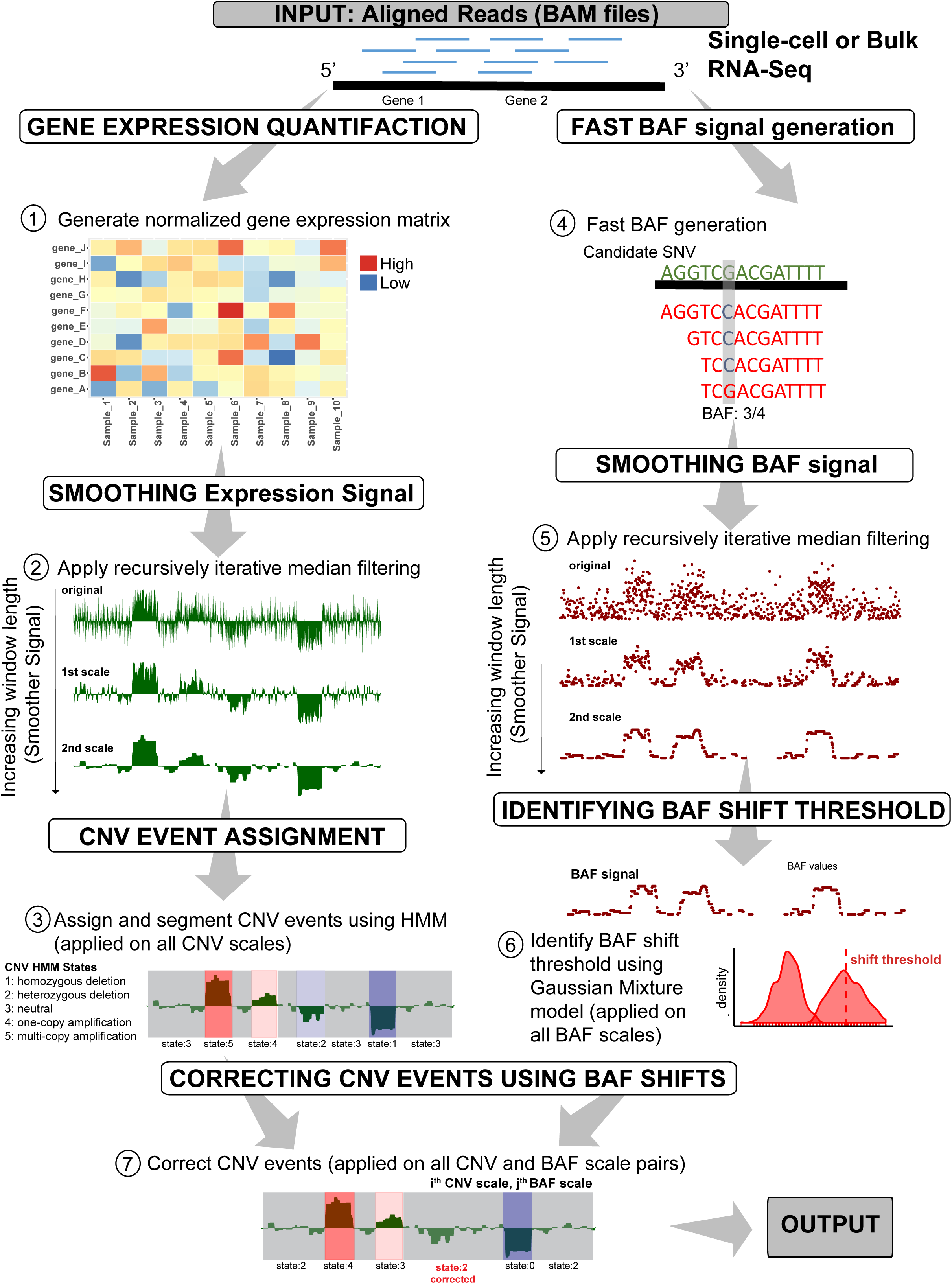
Flowchart of CaSpER algorithm. The CaSpER algorithm uses expression values and B-allele frequencies (BAF) from RNA-Seq reads to estimate CNV events. Normalized gene expression matrix is generated (Step1). Expression signal is smoothed by applying recursive iterative median filtering. Three scale resolution of the expression signal is computed. (Step2). For the smoothed signal at each scale, HMM is used to assign CNV states to regions and segment the signal into regions of similar copy number states (Step 3). Five CNV states are used in HMM model; 1: homozygous deletion, 2: heterozygous deletion, 3: neutral, 4: one-copy amplification, 5: multi-copy amplification. BAF information incorporated to the segmented CNV events. BAF information is extracted from mapped RNA-Seq reads using an optimized BAF generation algorithm (Step 4). BAF signal is smoothed by applying recursive iterative median filtering. Three scale resolution of the allele-based frequency signal is computed (Step 5). BAF shift threshold is estimated using a Gaussian mixture (Step 6). CNV events are corrected using BAF shifts and final CNV correction is applied to all the CNV and BAF scale pair combinations (Step 7).

CaSpER incorporates the allelic shift (BAF) information as a separate evidence of CNVs in addition to the CNVs identified from the multiscale analysis of genome-wide expression signal profile. BAF information is extracted directly from the mapped RNA-Seq reads using an optimized BAF generation algorithm (see Methods for details). Unlike other methods, BAF generation does not rely on an existing set of variant calls and this considerably speeds up the process of estimating the BAF signal. Similar to expression signal, BAF signal is also smoothed using multiscale decomposition. CaSpER smoothes BAF signal at multiple window length scales where window length is increased between consecutive scales. A CNV event with states 2 (heterozygous deletion) and 4 (one-copy amplification) as correctly identified if the CNV state is accompanied by BAF shift. BAF shifts are detected by thresholding the smoothed BAF signal profiles where the threshold is estimated by pooling BAF information across all samples and fitting a Gaussian Mixture Model on the distribution of the smoothed BAF values for the selected single nucleotide variations (SNVs) in segmented regions. CaSpER performs a pairwise comparison of all scales from BAF and expression signals to ensure a coherent set of CNV calls are detected from the multiscale decomposition. The final CNV event calls for all the CNV and BAF scale pair combinations are stored as the output from CaSpER’s CNV calling steps (Figure 1).

CaSpER outputs detailed focal CNV calls for all CNV and BAF scale pairs. Moreover, it outputs the large-scale CNV calls that are commonly seen in all scale pairs. For single-cell RNA sequencing studies, CaSpER infers CNV based clonal evolution depicted as a phylogenetic tree and summarizes mutually exclusive and co-occurring CNV events using graph-based visualization (Figure 2).

**Figure 2.**
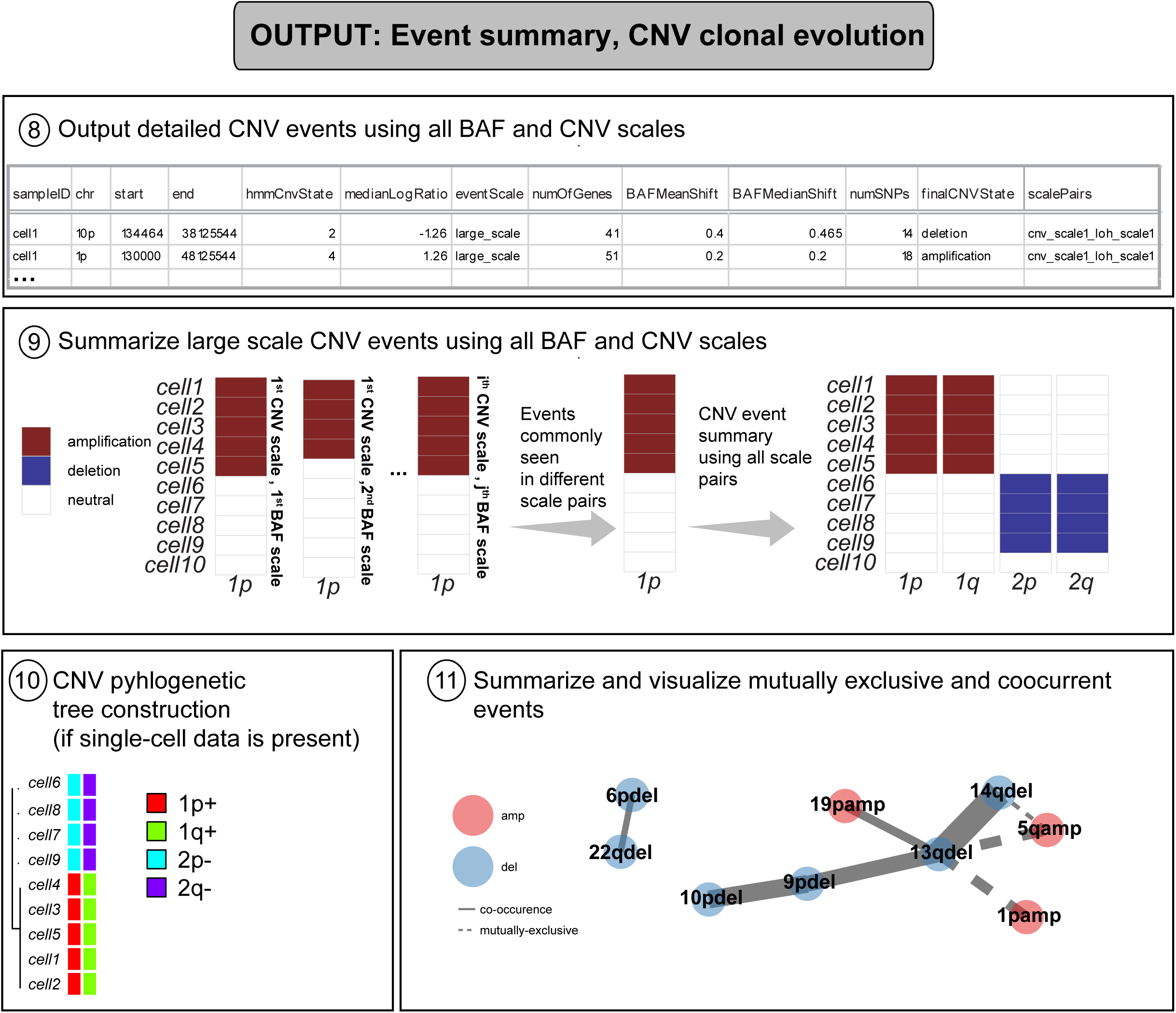
CaSpER algorithm outputs detailed CNV event calls for all CNV and BAF scale pairs (Step8). Large-scale CNV events that are commonly seen in all scale pairs are reported (Step 9). For single-cell RNA sequencing studies, CaSpER infers CNV based clonal evolution depicted as a phylogenetic tree (Step10). Mutually exclusive and co-occurring CNV events are summarized using graph-based visualization (Step11).

### Evaluation of the accuracy of CNV events detected from bulk RNA-Sequencing using genotyping array

We validated CaSpER algorithm on publicly available TCGA-GBM (n=171) and another separate meningioma cancer study (n=17), where both bulk RNA sequencing and genotyping data is available^24,25^. We explain below the outputs and accuracy of CaSpER on these datasets. For TCGA-GBM dataset, expression values of all the genes are quantified across all TCGA-GBM samples. The recursive median filtering effectively removes the fluctuations in the genome-wide expression introduced by noise and fluctuations in expression (Figure 3A). For the smoothed signal at each scale, we applied HMM to assign CNV states to segmented regions (Figure 3B). Simultaneously, BAF signal is extracted from RNA-Seq bam files using our fast BAF generation method and smoothed using recursive median filtering. BAF shift threshold is estimated by fitting Gaussian mixture model (GMM). GMM identified three classes of BAF shift groups where the first group corresponds to no shift regions whereas the second and the third group corresponds to BAF shift regions with loss or amplification events. (Figure 3C).

**Figure 3.**
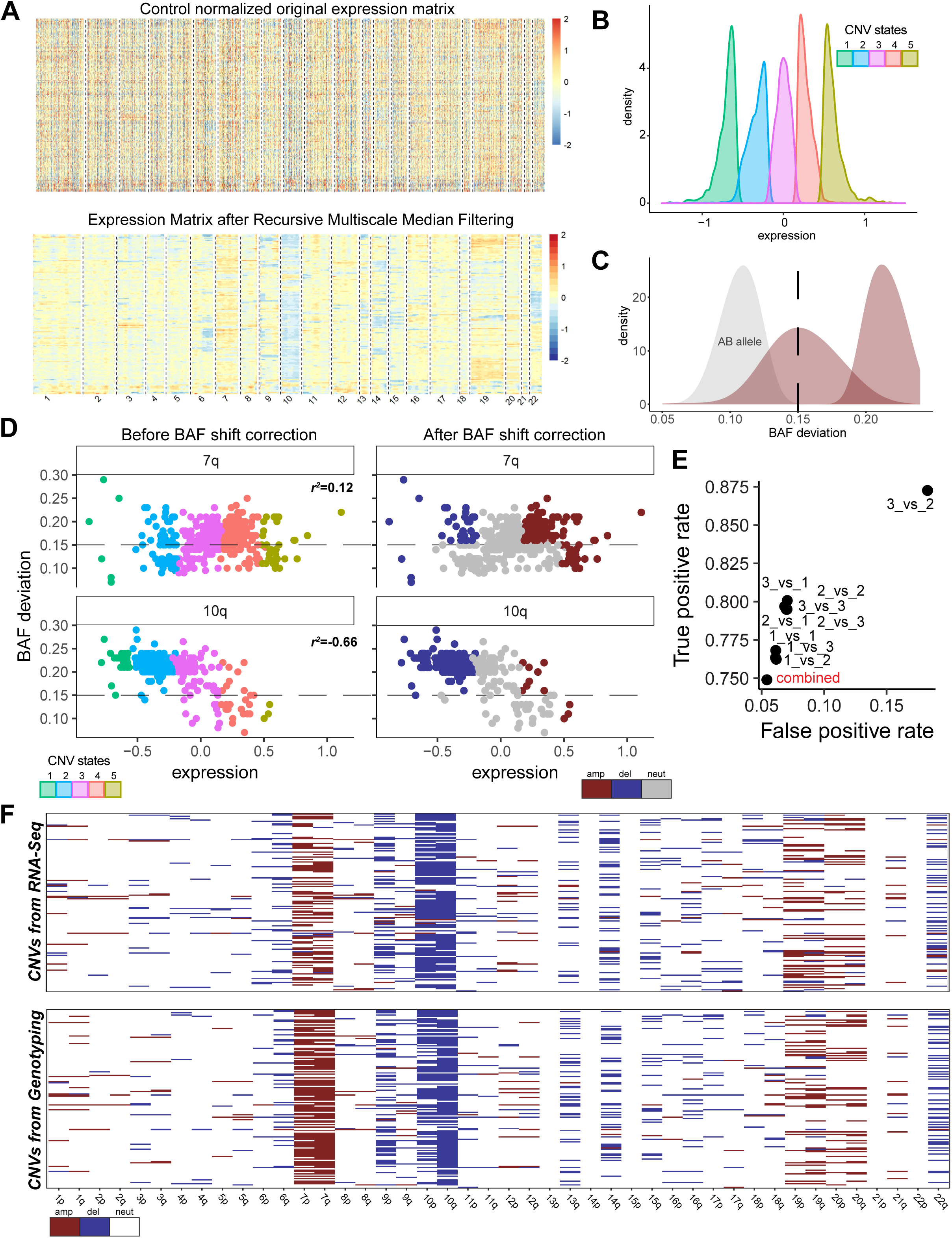
CaSpER algorithm applied to bulk TCGA-GBM RNA-Seq dataset. **A.** Heatmap of normalized expression values of all the genes across all TCGA-GBM samples (~170 samples) is shown in top panel. The smoothed expression signal by recursive iterative median filtering is shown on the bottom panel. **B.** For the smoothed signal at each scale, HMM is applied to assign CNV states to segmented regions. **C.** BAF shift threshold is estimated by fitting Gaussian mixture model (GMM). GMM identified three classes of BAF shift groups where the first group corresponds to no shift regions whereas the second and the third group corresponds to BAF shift regions with loss or amplification events. BAF shift threshold is the median of the BAF values in the second group, which is calculated to be 0.15. **D.** The correlation of expression values and BAF values in recurrently amplified chromosome 7q arm and deleted chromosome 10q arm is plotted (chr7p, r^2^=0.12, P=0.007; chr10q, r^2^=−0.66, P<2.2E^−16^). Regions with CNV states 2 and 4 that are below the BAF shift threshold (0.15) are corrected to be neutral. The color codes are explained on the bottom. **E.** The false positive and true positive rates of different CNV and BAF scale pairs is plotted. Summarized large-scale event calls have a lower false positive rate. **F.** Heatmap of large-scale CNV events identified from RNA-Seq is shown in the top panel whereas the heatmap of large-scale CNV events identified from genotyping is shown in bottom panel. The color codes are explained on the bottom.

We also investigated the correlation of expression values and BAF values in recurrently amplified chromosome 7q arm and deleted chromosome 10q arm. We observed a significant correlation between BAF and expression values (chr7p, r^2^=0.12, P=0.007; chr10q, r^2^=−0.66, P<2.2E^−16^) (Figure 3D). Regions with CNV states 2 (heterozygous deletion) and 4 (one-copy gain) that are below the BAF shift threshold are corrected to be neutral (Figure 3D).

We next used the CNV calls identified from genotyping arrays to measure the accuracy of CaSpER. For each sample, we identified the large-scale deletions and amplifications from genotyping array, where the large-scale event is defined as more than 1/3 of the chromosome arm. We next calculated the false positive and true positive rates (See Methods Section) of different CNV and BAF scale pairs. Summarizing the large-scale CNV events that are commonly seen in all scale pairs lowers our false positive rate (FPR) to 5%. This points out the importance of studying the CNV events at multiple scales. By using a genotyping array (n=160) as a gold standard, after summarizing CNV events using all scale pairs, CaSpER achieves 62% true positive rate (TPR) and 1% FPR for detecting amplification events and 84% TPR and 3% FPR for detecting deletion events (Figure 3E-F).

We also validated CaSpER on a publicly available meningioma dataset^24^. We first quantified the expression values of all the genes across all meningioma samples. After quantifying expression values, recursive median filtering is applied to eliminate noise from the signal. Heatmap of smoothed data clearly shows chromosome arm size deletion events (Figure 4A). HMM is applied to the smoothed signal at each scale to assign CNV states to segmented regions (Figure 4B). The smoothed signal distribution shows that data does not contain many amplification events (Figure 4B). Concurrently, BAF signal is calculated from aligned RNA-Seq reads using our fast BAF generation method. Similar to expression signal, BAF signal is also smoothed using recursive median filtering with iterative scale lengths (Figure 4C). Smoothed BAF signal shows shifts in chromosomes with deletion events (Figure 4C). For each scale, GMM is fitted to the BAF values and identified two sets of events. First set of copy number events contains BAF values without shift whereas the second set contains BAF values with the shift. (Figure 4D). We observed negative correlation between BAF and expression values at chromosomes that are recurrently deleted (chr1p, r^2^=−0.43 P =0.0008; chr6q r^2^=−0.54 P =0.0003; chr22q r^2^=−0.14 P =0.43) (Figure 4E). We validated the accuracy of CaSpER using an existing genotyping array for the same dataset as the gold standard. After summarizing CNV events using all scale pairs, CaSpER achieves 95% TPR and 0.3% FPR for detecting deletion events (Figure 4F).

**Figure 4.**
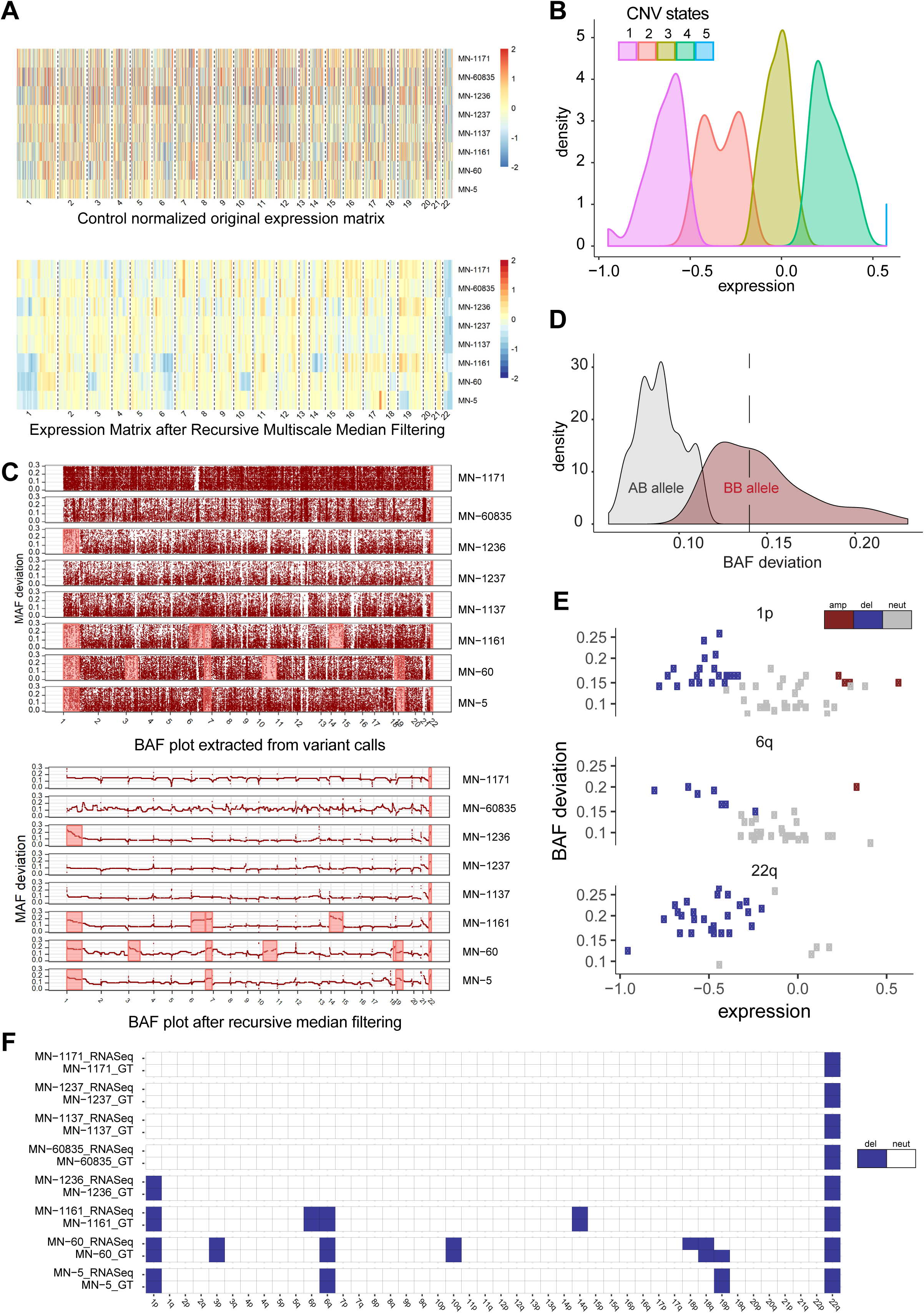
CaSpER algorithm applied to bulk meningioma RNA-Seq dataset. **A.** Heatmap of normalized expression values of all the genes across all samples (n=8 samples) is shown in top panel. The smoothed expression signal by recursive iterative median filtering is shown on the bottom panel. **B.** For the smoothed expression signal at each scale, HMM is applied to assign CNV states to segmented regions. **C.** Smoothed BAF signal is plotted and shows shifts in chromosomes with deletion events. **D.** BAF shift threshold is estimated by fitting Gaussian mixture model (GMM). GMM identified two classes of BAF shift groups where the first group corresponds to no shift regions whereas the second group corresponds to BAF shift regions with loss or amplification events. BAF shift threshold is the median of the BAF values in the second group, which is calculated to be 0.14 **E.** The correlation of expression values and BAF values in recurrently deleted chromosomes are plotted (chr1p, r^2^=−0.43 P =0.0008; chr6q r^2^=−0.54 P =0.0003; chr22q r^2^=−0.14 P =0.43). **F.** Heatmap of large-scale CNV events identified from RNA-Seq and genotyping is shown in the plot. The color codes are explained on the right.

We observed a slight difference in deletion TPR rates, between meningioma and TCGA-GBM datasets (95% TPR meningioma vs 84% TPR GBM). This stems from the fact that meningioma tumors exhibit less intratumor heterogeneity and have lower clonality rates compared to GBM tumors. Thus, lower clonality rates in meningiomas lead to better deletion CNV event identification. Similarly, high clonality rates in GBM tumors lower the detection accuracy of low-level amplification events, which then lead to low amplification TPR rate.

### Inference of subclonal CNV architecture in single-cell RNA-Sequencing data

We next used CaSpER to infer subclonal CNV architecture from single-cell Glioblastoma Multiforme (GBM) RNA sequencing data^8^. Single-cell GBM RNA-Seq data contains 430 single cells extracted from five patient samples. The smoothed expression signal in Figure 5A shows that chromosome 7 amplification and chromosome 10 deletion is recurrent across GBM samples. We observed that the BAF signal is much more stable and accurate when the data from all the cells are pooled. Therefore, for BAF signal generation step, we pooled the single cell reads from the same patient together. We extracted the BAF signal from the pooled patient specific reads instead of single cell reads since the BAF signal extracted from one single cell is very sparse and is not informative. The smoothed patient specific BAF signal shows shifts in chromosome 7 and 10 (Figure 5B). We can also detect a subclonal shift in chromosome 14 for patient MGH31 (Figure 5B). We next used the large-scale CNV events summarization to identify the common events in all scale pairs (Figure 5C). Interestingly, MGH31 consists of two mutually exclusive subclones where one subclone contains chromosome 5q amplification whereas the other subclone contains chromosome 14q deletion (Figure 5C). Additionally, one subclone contains 1p amplification and the other subclone contains 13q deletion, which has not been reported previously. We then evaluated the phylogenetic tree based visualization of the inferred subclonal CNV architecture as reported by CaSpER for all the five patients using large-scale CNV events. For MGH31, tree separated cells harboring 1p and 5q amplification from cells harboring 13q and 14q deletion (Figure 5D). CaSpER also reports the mutually exclusive and co-occurring CNV events and plots the significant events as a graph. This is useful for visually inspecting the co-occurring and mutually exclusive events that may be otherwise hard to visualize. Similarly, from the graph, we can clearly see the mutually exclusive 1p:13q and 5q:13q, 5q:14q event pairs for patient MGH31 (Figure 5E). Moreover, we detected novel mutually exclusive 8q:20p, 5q:19p event pairs for patient MGH28, 6p:7p event pair for patient MGH30 and 4p:10p event pair for patient MGH29 that was not reported in previous publications (Figure 5F, Supplementary Figure S1). We next identified the gene expression signatures of each of the mutually exclusive clones and performed GO enrichment analysis (Supplementary Data 1). For 5q:14q event pair in patient MGH31, we discovered *GFPT2* gene to be highly expressed in 5q amplified clone. Previous study has discovered that higher expression of *GFPT2* is linked with poor survival and identified *GFPT2* gene to be a potential target for therapeutic inhibition^26^. For 5q:19q event pair in patient MGH28, we discovered *NOS2* gene to be highly expressed in 19q deletion clone. It has been previously demonstrated that *NOS2* gene is expressed in glioma stem cells and high expression of *NOS2* is correlated with decreased survival^27^. Gene expression signatures of each of the mutually exclusive clones and GO enrichment analysis are reported in Supplementary Data 1. These findings demonstrate that the diverse set of visualizations and results that CaSpER generates enables studying the single cell RNA-sequencing data comprehensively over many different length scales. In addition, these results show that RNA-seq datasets contain much information about CNV and transcriptional architecture of the samples, which may otherwise be left out when standard analysis pipelines are used.

**Figure 5.**
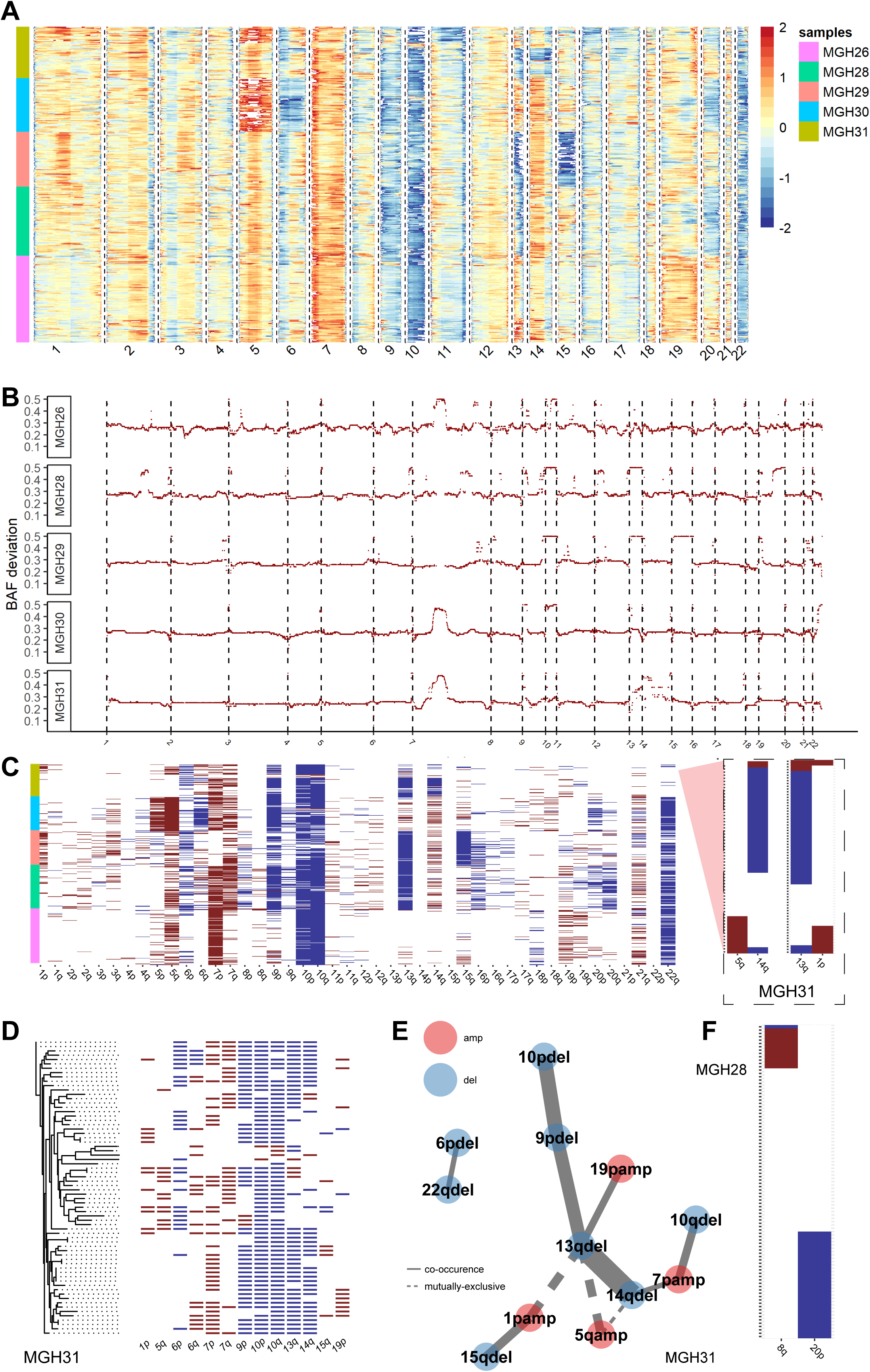
CaSpER algorithm applied to single-cell GBM RNA-Seq dataset. **A.** Heatmap of smoothed expression signal of all the genes across all samples is shown in top panel. The color codes are explained on the right. **B.** Smoothed BAF signal from the pooled patient specific reads is shown in the plot. The smoothed patient specific BAF signal shows shifts in deleted and amplified chromosomes. **C.** The heatmap of summarized large-scale CNV events using the common events in all scale pairs is plotted. MGH31 consists of two mutually exclusive sub-clones where one sub-clone contains chromosome 5q amplification whereas the other sub-clone contains chromosome 14q deletion. Additionally, one sub-clone contains 1p amplification and the other sub-clone contains 13q deletion. **D.** Inferred sub-clonal CNV architecture is shown as a phylogenetic tree for all the five patients using large-scale CNV events. For MGH31, tree separated cells harboring 1p and 5q amplification from cells harboring 13q and 14q deletion. **E.** Mutually exclusive and co-occurring CNV events are plotted as a graph. Red colored events are amplified whereas blue colored events are deleted. The solid lines represent co-occurring events, whereas dashed lines represent mutually exclusive events. Edge width increases with event significance. The mutually exclusive 1p:13q and 5q:13q, 5q:14q event pairs for patient MGH31 is significant. **F.** Novel mutually exclusive 8q amplification and 20p deletion CNV event in patient MGH28 is plotted.

### Identification of scale-specific CNV regions (SSCNVs) in bulk RNA and single-cell RNA-Sequencing data

CaSpER identifies CNV regions at each scale, which yields scale-specific CNV regions (SSCNVs). In general, SSCNVs at lower scale correspond to more focal CNV events compared to SSCNVs at higher scale, which represents broader CNV events. Figure 6 shows the scale length characteristics of different CNV events of TCGA bulk RNA-Sequencing data. Focal amplification of the *PDGFRA* gene is identified using small-scale lengths whereas broad chromosome arm level deletion in chromosome 22 is identified using a higher scale length (Figure 6A-B).

**Figure 6.**
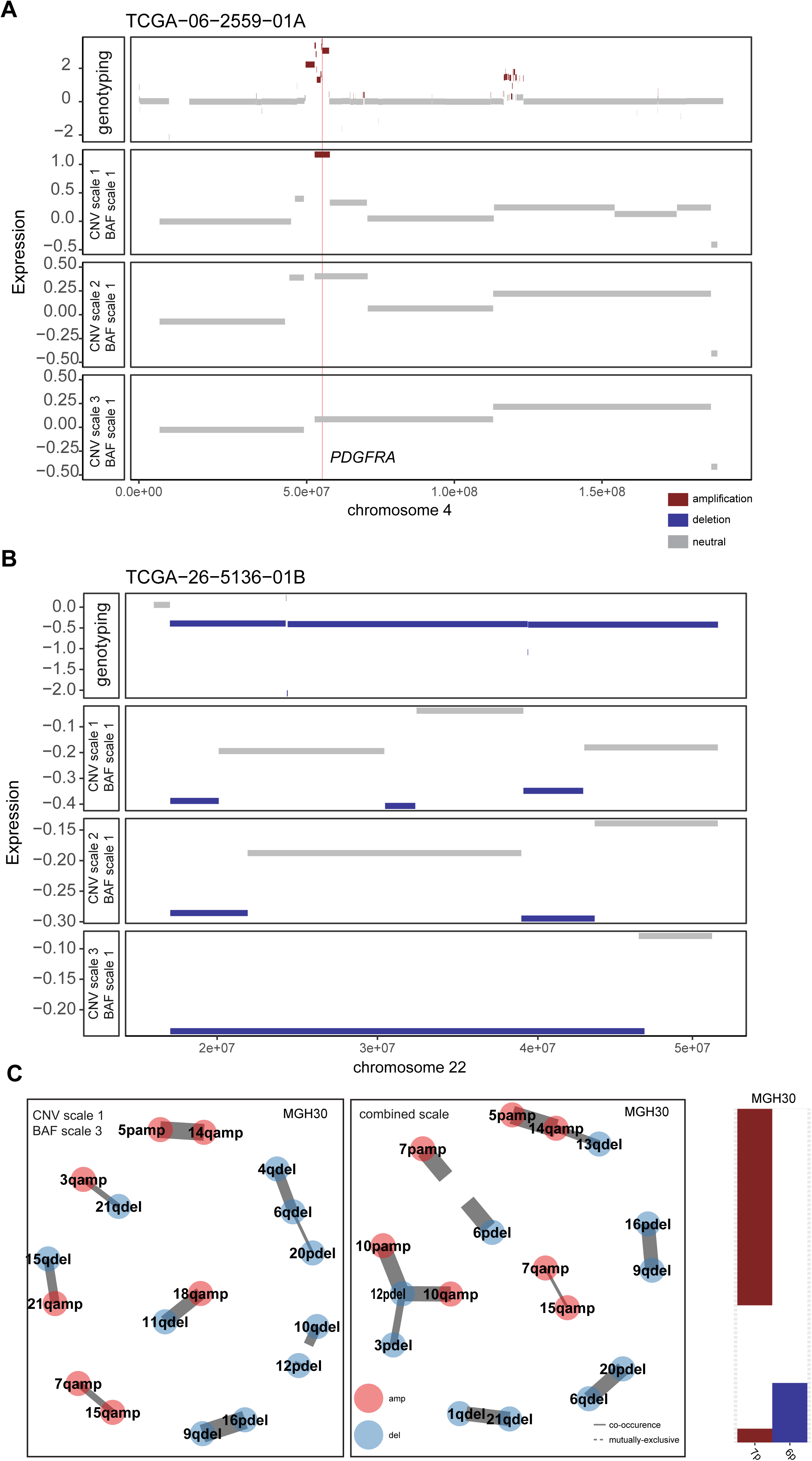
CaSpER identifies scale-specific CNV regions (SSCNVs). **A.** PGFRA focal amplification is identified using lower scale length. **B.** Broad chromosome arm level deletion in chromosome 22 is identified using a higher scale length. **C.** Scale-specific sub-clonal events for patient MGH30 in single-cell RNA sequencing data is plotted. Mutually exclusive 7p:6p event pair is detected after summarizing large-scale CNV events in all scales.

We next investigated scale specific sub-clonal events in single-cell RNA sequencing data (Figure 6C). We observed that different scale pairs are capable of identifying different mutual-exclusive or co-occurring CNV events. One challenge in the multiscale analysis is a summarization of a large number of data from multiple scales. To get around this, CaSpER analyzes and automatically generates a visualization of the significant mutual exclusive or co-occurring CNV events as heatmaps and graphs. These visualizations are especially useful for visually confirming the significance of the results. Figure 6C shows that mutually exclusive 7p:6p event pair is detected after summarizing CNV events using all scale pairs (Figure 6C).

### Identification of BAF shifts in bulk and single-cell RNA sequencing data

We expect that regions with amplification or deletion to be mostly accompanied with loss of heterozygosity (LOH) and harbor BAF shift except for homozygous deletion or amplification regions. In most of the cases, we do not have matched normal RNA-Seq samples, therefore; we predict this BAF shift (LOH) regions only from tumor RNA-Seq samples.

We compared our BAF signal with the BAF signal generated from GATK best practices workflow for RNA-Seq variant calling. We did not observe difference between our BAF signal and the BAF signal generated using GATK tool (Supplementary Figure S2A-B)^14^.

In bulk tumor RNA Sequencing, the presence of a normal population in the tumor, also known as impurity or admixture of normal cells in a tumor sample, and sub-clonal events may add complex noise into the BAF signal. The SNP sites that are around the band of 1 are not considered since these sites are most likely to be homozygous in normal samples. Because of admixture rate, it is unlikely to see heterozygous sites in normal to have BAF value around 1. If we also consider the BAF values more than 0.8, not only the original BAF signal but also the smoothed BAF signal is very noisy (Supplementary Figure S2B-C). Another important factor in generating reliable smooth BAF signal is to filter out reads with low mapping quality (Supplementary Figure S2B, 2D).

In single-cell RNA Sequencing, we pool single cells belonging to the same patient since the BAF signal for one single cell is not informative (Supplementary Figure S3). In single cell RNA-Sequencing, we can pool only the tumor single cells of the patient and eliminate all other cell types such as immune cells. Since we do not have the admixture rate issue in single cell RNA-Sequencing data, the SNPs with BAF value more than 0.2 are considered (Supplementary Figure S4).

## Discussion

We presented a novel algorithm, CaSpER, for identification, visualization, and integrative analysis of focal and large-scale CNV events in multiscale resolution using either bulk or single-cell RNA sequencing data. We demonstrated that CaSpER performs well in identifying CNV events using both single-cell and bulk RNA-Sequencing data. We presented several examples where CaSpER can effectively complement the existing set of RNA-Seq analysis tools and can also identify new insight into the analysis of the clonal architecture of cancer genomics datasets. CaSpER can be used to efficiently do a comprehensive characterization of RNA-Sequencing datasets to increase their utility beyond transcriptional profiling.

There are several aspects of CaSpER that we would like to point out. First, CaSpER combines genomewide allelic shift signal, which measures the loss-of-heterozygosity at a nucleotide resolution, and expression signal to accurately estimate CNV events. While doing this, CaSpER utilizes a novel method to generate allelic shift signal profile directly from mapped reads, without the need for variant calls. This is very useful for two reasons: First, the variant calling (SNVs and indels) from mapped reads requires high computational resources and long compute times. This means that before calling CNVs, it is necessary to complete the arduous process of variant calling. CaSpER lifts this restriction by computing the allele shift profile directly from the mapped reads. Second, the power of variant detection can be affected by the CNV events. This is especially important in cancer sequencing experiments where CNVs can span very long genomic regions and may affect the accuracy of variant calling. Therefore, identifying CNVs before calling SNVs can give very useful information for correct identification of SNVs. Although we did not explore this thoroughly in this paper, several previous studies have demonstrated this^28^.

Another novel aspect of CaSpER is the analysis of CNV events in multiscale resolution. With the diverse length characteristics of CNV events, we believe that it is very important to be able to analyze CNV events in multiple length scales. CaSpER makes available the multiscale smoothed genomewide expression signal and allelic shift signal profiles and the CNV calls that can be used for downstream analysis and visualization. For smoothing, CaSpER utilizes a non-linear median based filtering of RNA-Sequencing expression and allele-frequency signal. It is worth noting that median filtering preserves the edges of the signal much better compared to kernel-based linear filters^23^. We also demonstrated that the signal profiles that are smoothed at multiple scales are useful for visualization of the copy number events that are detectable from RNA-seq datasets. In addition to identifying CNV events, CaSpER also visualizes and performs integrative analysis of CNV events such as inferring clonal evolution, discovering mutual-exclusive and co-occurring CNV events and identifying gene expression signatures of the identified clones.

In this paper, we have focused on detection of CNV events from RNA-Seq datasets. This is because RNA-sequencing is becoming an everyday tool in research and clinical settings. Also, the number of RNA-sequencing datasets is increasing comparably with the whole genome and whole exome sequencing datasets. Although we have focused only on RNA-sequencing data, the analysis framework that CaSpER utilizes can be extended to other functional genomics datasets such as ChIP-Sequencing, which are currently not performed as often as RNA-sequencing. We hypothesize that the CNV architecture can be reliably detected using these functional genomics datasets jointly. As more cancer epigenomics datasets are generated, CaSpER can be tuned to analyze data from these assays for detection of copy number and LOH events.

In our analyses, CaSpER discovered scale-specific CNV regions in TCGA bulk RNA-sequencing data, which represent both focal and broad CNV events and we showed the utility of these events in the analysis of scale-specific co-occurrence and mutual exclusivity of the CNV events. Analyzing single-cell GBM RNA Sequencing data using CaSpER, unraveled novel mutual exclusive and co-occurring CNV subclones. Gene expression signatures of the identified novel clones gave us insight about the phenotype of the clones such as invasiveness and survival. Moreover, we identified novel potential therapeutic targets for the clones. In conclusion, our study demonstrates the significance and feasibility of CNV calling using either single or bulk RNA sequencing data.

## Methods

### Bulk and Single-Cell RNA-Sequencing expression quantification

Yale meningioma bulk RNA-Sequencing data reads were aligned using STAR^29^. Expression level quantification was performed using DESeq2 R package^30^. TCGA-GBM bulk RNA-Sequencing normalized expression matrix were downloaded using TCGABiolinks R package^31^. The corresponding bam files were downloaded from GDC data portal^32^.

Single-cell RNA Sequencing data reads were aligned with Hisat2 using ENCODE V28 transcriptome annotation^33^. We pooled single cells from the same patient into a single bam file using bamtools merge function^34^. The aligned bam files were later used for allele-based frequency signal calculation. We used the normalized expression matrix provided in the paper.

### Generation of the allele-based frequency signal from RNA-Seq BAM files

We generate allele-based frequency signal from RNA-Seq bam files using our in-house written C++ code. Our method takes a bam file as an input and outputs allelic content estimation through fast SNP calling. We first perform pileup where we summarize the base calls of aligned reads to a reference sequence. For each SNP, we report the total count of reads supporting non-reference and reference nucleotide after applying the following filters: (1) reads should have mapping quality of at least 50, (2) minimum number of total reads per each SNP position should be 20 (3) minimum number of total reads supporting SNP should be 4. In bulk tumor RNA-Seq, we considered SNPs that are most likely to be heterozygous with a BAF value more than 0.2 and less than 0.8 whereas in single-cell RNA-Seq BAF value more than 0.2 are considered. Our fast BAF generation method speeds up the process of estimating BAF shift regions compared to using GATK for calling variants from RNA-Seq data. After generating the allelic content for each SNP, we next apply recursive median filtering to remove noise from the signal. As explained previously in the Results section, filtering the reads according to mapping quality is very critical for correctly estimating BAF shift regions. In addition to estimating BAF shift regions, our method is also very useful in identifying allele-specific expression.

### Multiscale resolution of expression by median filtering

Unlike linear filtering methods, median filtering preserves the edges while removing noise in smooth regions of signal^23^. CaSpER uses recursive median filtering for removing noise from expression and allele-based frequency signal.

Let
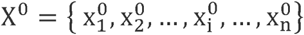
be the expression signal vector, where
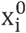
is original signal value at iteration 0 in position i. Given the window length l, at scale s median filtering can be formulated as:

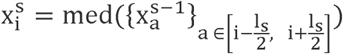

where
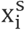
is the i^th^ value of the median filtered expression signal at scale s. In
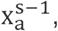
a is defined as smoothing region of each i formulated as
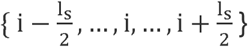
and the input expression signal x^s−1^ is the smoothed expression signal in the previous iteration, s − 1.

Similar to expression signal we also apply recursive median filtering to an allele-based frequency signal. Let
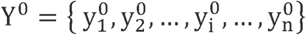
be the expression signal vector, where y is the original allele-based frequency signal value at iteration 0 in position i. Given the window length l, at scale s median filtering can be formulated as:

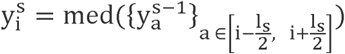

where
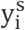
is the i^th^ value of the median filtered allele-based frequency signal at scale s. In
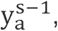
a is defined as smoothing region of each i formulated as
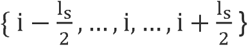
and the input y^s−1^ is the smoothed allele-based frequency signal in the previous iteration, s − 1

CaSpER uses *filter* function in *signal* R package for median filtering implementation.

### Gaussian mixture models

CaSpER models the allele-based frequencies as a mixture of Gaussian distributions for identification and classification of genotype clusters. For example, in a normal chromosomal region with 2 copies, we expect to observe three BAF genotype clusters represented as AA, AB, and BB whereas, in heterozygous deletions, we expect to observe two clusters which can be represented as A and B.

Let X = {x_1_, x_2_, …, x_i_, …, x_n_} be the allele-based frequency signal vector, where x_i_ is the signal value at position i. The distribution of every value is specified by a probability density function through a finite mixture model of G classes:

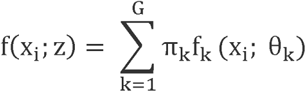

where z = {π_1_, ….,π_G−1_, θ_1_, … θ_G_} is the parameters of the mixture model and f_k_ (x_i_; θ_k_ is the kth component density, which assumes to follow Gaussian distribution f_k_ (x_i_; θ_k_) ~ N(µ_k_,σ_k_). {π_1_, …., π_G−1_} is the vector of probabilities, non-negative values which sum to 1, known as the mixing proportions. Mixing proportions, π, follows a multinomial distribution.

The model z parameters are estimated by maximizing log-likelihood function via the EM algorithm. The log-likelihood function is formulated as:

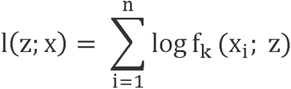

The number of classes, G, are estimated using the Bayesian Information Criteria (BIC). The class with the lowest mean value corresponds to alleles without any BAF shift. We choose the class with second lowest mean value, called as ‘*class 2*’ to identify the BAF shift threshold. In bulk sequencing data, we set the BAF shift threshold to mean allele-based frequency signal in ‘*class 2*’. In single-cell RNA sequencing data, we set the BAF shift threshold to minimum allele-based frequency signal in ‘*class 2*’. CaSpER uses mclust R package for Gaussian mixture model (GMM) implementation^35^.

### Hidden Markov model (HMM)

CaSpER uses a modified version of HMMCopy R package for HMM implementation to 1) segment the copy number profile in regions predicted to be generated by the same copy number event and 2) predict the copy number variation event for each segment. In HMM, we use a hierarchical Bayesian model where the posterior estimates are calculated using an exact likelihood function.

Our HMM model contains 5 CNV states, where the states represent homozygous deletion, heterozygous deletion, neutral, one-copy gain and multiple-copy gain. The initial transition matrix is defined as:

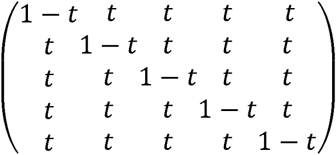

where t is equal to 1e-07. We estimated the mean parameter of the emission probabilities by pooling all the chromosomes across all the samples. We believe that using more data by pooling all the samples would give better initial HMM parameter estimates.

### Calculating mutual-exclusive and co-occurring CNV events and inference of clonal phylogenetic CNV tree

We calculated the distance between cells using the Jaccard distance metric. We next used this distance matrix to build the phylogenetic tree of the CNV events. A phylogenetic tree is constructed using Fitch–Margoliash method implemented in the Rfitch R package. We finally plotted the tree using phydataplot function in ape R package. The co-occurrence and mutual exclusivity of CNV events were assessed using one-sided Fisher’s exact test.

### Identifying gene expression signatures and enrichment analysis

Differentially expressed genes were identified using an empirical Bayesian method ebayes implemented in limma R package^36^. Genes were considered differentially expressed with adjusted *P*-value<0.05. GO term enrichment analysis was performed using the GOStats R package^37^.

### Genotyping data

TCGA-GBM genotyping data was downloaded using TCGABiolinks R package^31^. CNV segments with mean log ratio value more than 0.3 were defined as amplification whereas segments with mean log ratio value less than 0.3 were defined as deletion. Large-scale chromosomal deletion or amplification was defined as affecting more than one-third of the chromosomal arm, whereas focal event deletion or amplification was defined as affecting less than one-third and more than one-tenth of the chromosomal arm with accompanying log ratio of signal intensities <−0.1 or >0.1 and B-allele frequencies (BAF) at heterozygous sites deviating from 0.5 by at least 0.05 units.

We analyzed meningioma genotyping data as previously described^24^. CNVs were detected by comparing the normalized signal intensity between a tumor and matched blood or a tumor and the average of all blood samples. Segmentation was performed on log intensity (R) ratios using DNACopy algorithm^38^.

### Validating Bulk RNA-Sequencing results using genotyping data

We assessed the performance of CaSpER by comparing the CNV calls identified from RNA-Seq with genotyping data. Thus, the true positive rate is the percentage of large-scale CNV events who are correctly identified by CaSpER while the false positive rate is the percentage of falsely rejected true CNV events.

## Acknowledgements

This study is partially funded by the NIH grants (R01GM123037, U01AR69393, U01CA166866) and STARs award.

